# OPTIMIZING SEGMENTATION IN OCCUPANCY MODELLING OF CAMERA-TRAP DATA

**DOI:** 10.1101/2024.06.11.598409

**Authors:** Monique de Jager, Marijke van Kuijk, Joeri A. Zwerts, Patrick A. Jansen

## Abstract

1. Accurate estimation of species’ abundances is a common challenge in conservation biology, particularly when abundances are compared in space or time. Occupancy modelling provides relative abundance estimates from camera-trapping data without the need for individual recognition. This requires segmentation of continuous records into a series of intervals with either detection or non-detection. While the segmentation method may have profound effects on the accuracy of occupancy modelling, no form of segmentation optimization is yet available.
2. We assessed how segmentation, defined by interval length and number, influences the accuracy of predictions by the Royle-Nichols occupancy model and how this relationship depends on species’ density, study duration, and the number of sampling points. We simulated capture data using an individual-based model in which we varied the species’ densities between study locations, and then fitted models using different segmentations. Using the simulation results, we developed a simple tool for choosing optimal segmentation and the best minimum number of intervals to use. To provide an example, we used the optimization tool on actual data from a camera-trapping study in Western Equatorial Africa and compared relative wildlife abundances between two forest management types.
3. We found that the optimum interval length for the Royle-Nichols occupancy model varied with species’ density, study duration, and the number of sampling points. By analyzing the empirical data, we found that optimal segmentation and minimum number of intervals differed substantially between species. Modelling with optimized, species-specific interval numbers and lengths yielded more conservative outcomes (i.e. fewer significant effects) than did modelling with fixed numbers and lengths. Furthermore, the choice of interval length can affect the direction of relationships.
4. Our results indicate that the interval length is by no means a parameter to be standardized at a given value but should be carefully chosen based on the properties of the data at hand. This study shows that the arbitrary segmentation that is commonly used in occupancy modelling may not be optimal. Our tool helps to optimize segmentation, increases the accuracy of relative abundance estimations, and thus facilitates the use of camera-trapping studies to evaluate conservation measures.

## 1. Introduction

Camera trapping is a widely used method for estimating abundance and monitoring changes in wildlife populations over time, contributing to wildlife conservation efforts and ecological research (O’Connell et al. 2011, Steenweg et al. 2016, Oliver et al. 2023). For species that cannot be individually identified on camera-trap images, a frequently used metric of abundance is occupancy: the proportion of area, patches or sites occupied by a species (MacKenzie et al. 2002, O’Connell and Bailey 2011). Classical occupancy modelling is typically used to determine the overall level of occupancy for a species in a site, or to compare the level of occupancy between different areas or habitats within a site (MacKenzie and Royle 2005).

The challenge that occupancy modelling solves is that of imperfect detection: species may be present at a sampling unit – e.g., a site – but nevertheless not captured. Occupancy models account for this by estimating the detection probability from multiple repeated surveys – i.e., ‘sampling occasions’ –yielding a series of ones and zeroes indicating detection and non-detection for each sampling occasion. Occupancy models relate the number of observations to the probability of detection and derive a probability of occupancy, which corresponds to the probability that at least one individual is available for detection at a site (Welsh et al. 2013).

In case of camera-trap surveys, repeated surveys are obtained by segmentation, i.e. subdividing the total sampling period of a camera deployment at a certain location into a number of time intervals with or without detections of the species of concern (e.g., Ahumada et al. 2013). Thus, selecting the number of repeated surveys that should be used (MacKenzie and Royle 2005), or, in case of camera trapping, the number of time intervals into which the detection record should be subdivided, is a key aspect of occupancy modelling. There have been several studies that evaluated the number of repeated surveys needed to detect differences and trends given different detection probabilities (MacKenzie et al. 2002, Stauffer et al. 2002, Tyre et al. 2003, Field et al. 2005), yielding rules of thumb. MacKenzie and Royle (2005), for example, recommend that sampling units should be surveyed a minimum of three times when detection probability is high (> 0.5).

The choice of interval length (the deployment duration divided by the number of intervals) has been more arbitrary, as continuous records can be segmented in many ways. The interval length varies widely among camera-trap studies, from as short as one day (e.g., Ahumada et al. 2011, Beaudrot et al. 2016) to as long as 30 days (e.g., Sunarto et al. 2015). The number of intervals chosen in the camera-trapping literature typically exceeds ten, and may even exceed 100 (Ahumada et al. 2011, Akkawi et al. 2020). Moreover, in multi-species studies, the common practice is to choose a single number of intervals and thus a single time-interval length for all species (Bowler et al. 2017, Van Kuijk et al. 2022). In a screening of the literature, we find that the choice of interval number and length is rarely justified explicitly, but most studies seem to aim for a large number of short intervals, as to retain more of the detections.

It is, however, plausible that the balance between interval length and interval number affects the occupancy model’s outcomes. Selecting short intervals (i.e. 1 day) may lead to zero inflation (Denes et al. 2015), while selecting larger intervals might result in an excess of ones in the occupancy data, leading to false conclusions about a species’ ‘occupancy’. In theory, there should be an optimum interval segmentation for modelling occupancy that depends on the species’ number of observations, which is in turn a function of actual abundance, detectability, number of sampling points, and study length. However, while there are clear evidence-based guidelines for optimization of the number of sampling points and the sampling duration (e.g., Gálvez et al. 2016, Kays et al. 2020), we have not encountered any study in which segmentation of camera-trap data for occupancy modelling was based on a form of optimization.

We studied how segmentation of camera-trap data influences the accuracy of occupancy models in scenarios with varying species density, study duration, and number of sampling points. We do this with the Royle-Nichols (RN) occupancy model (Royle and Nichols 2003), which deals well with heterogeneity in the detection probability that characterizes camera-trap data (Tobler et al. 2015). We first simulate capture data using an individual-based model in which we varied the species’ average density as well as differences in density between study locations. We then assess how accurate the actual density is estimated by fitted models that use different segmentations. With the results, we present a simple tool (i.e. R-script) for optimizing segmentation based on study length and the number of observations. Finally, we demonstrate the use of the tool by applying it to data from a camera-trapping study in Western Equatorial Africa (Republic of Congo and Gabon).

## 2. Effects of segmentation on accuracy

### 2.1. Data simulation

To assess the effect of interval length on model accuracy, we first generated semi-realistic camera-trapping data, using a simple, spatially explicit individual-based model (IBM), similar to the approach in Neilson et al. (2018). We simulated data for 19 virtual locations with known animal densities ranging from 0.04 to 3.2 individuals km^-2^. For the 15 highest densities, individuals moved randomly within a 10 x 10 km area consisting of 2,000 x 2,000 patches (i.e., one cell represented a 5 x 5 m area). For the four lowest densities, we used a larger area to increase the number of individuals in the simulation (35 x 35 km, consisting of 25,000 x 25,000 patches; Fig. S1). Of all the patches, 25 represented camera traps, which were equidistantly placed (5 x 5 m, with a 2 km distance between subsequent camera traps) in a 10 x 10 km area (Fig. S1). For a period of 365 days, *n* individuals moved across the area, using steps of three patches per minute and turning in a random direction after each step. Movement directions were drawn from a uniform distribution (0 < α ≤ 2π). During nighttime, movement stopped for 11 consecutive hours. If an individual moved outside the simulation area, it re-appeared on the other side (i.e. periodic boundary conditions). Whenever an individual was located at a camera-trapping patch, the time-step and camera number were logged. Due to the stochastic nature of the simulations, each density was simulated 10 times.

Camera-trap datasets for testing were obtained by subsampling each of the 19 simulated datasets and creating detection histories using interval lengths ranging from 1 to 110 days. The RN model depends on an input matrix that contains presence/absence (1/0) data per camera (= row) and time interval number (= column). Because of the daily activity cycle that most animals have, the interval length used in the RN model must be a multiple of one day. To examine the effect of study duration, we generated 16 different durations by cutting off the collected simulation data at *T* = 10, 15, 20, 25, 30, 40, 50, 60, 75, 90, 110, 135, 165, 200, 245, and 300 days. Subsampling was also used to generate datasets with different numbers of camera traps. Then, to assess the power of occupancy models detecting differences in density, we generated all possible combinations of pairs of simulated locations (where density A < density B) and fitted a RN model. In the RN model used on the modelled data, detection probability (*r*) of a species did not differ between locations and the abundance estimate (*λ*_*i*_) of a species was calculated per location *i*:

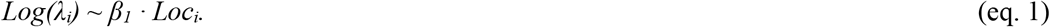

We recorded whether the RN model estimated a significantly higher abundance in location B than in location A (i.e., z > 1.96, or p < 0.05). Data simulation and statistical analyses were performed in R version 4.1.2 (R Core Team 2022) using the *occuRN* function in package unmarked for the RN model analyses (Fiske and Chandler 2011).

### 2.2 Model accuracy

Whether the RN model provides correct significant results depended on the modeled densities, the study duration, segmentation (i.e. time interval length), and number of cameras used (Fig. 1). When densities were too similar, most of the RN models were incorrectly showing that there was no significant difference in densities between locations, as defined by the occuRN function. For larger density differences between locations A and B, the percentage of RN models that provided correct results increased with study duration. The effect of time interval length was smaller, but still significant (p << 0.001), with a larger percentage of correct results for smaller interval lengths. The number of cameras used in the study positively affected the percentage of correct results (p << 0.001).

**Figure 1:**
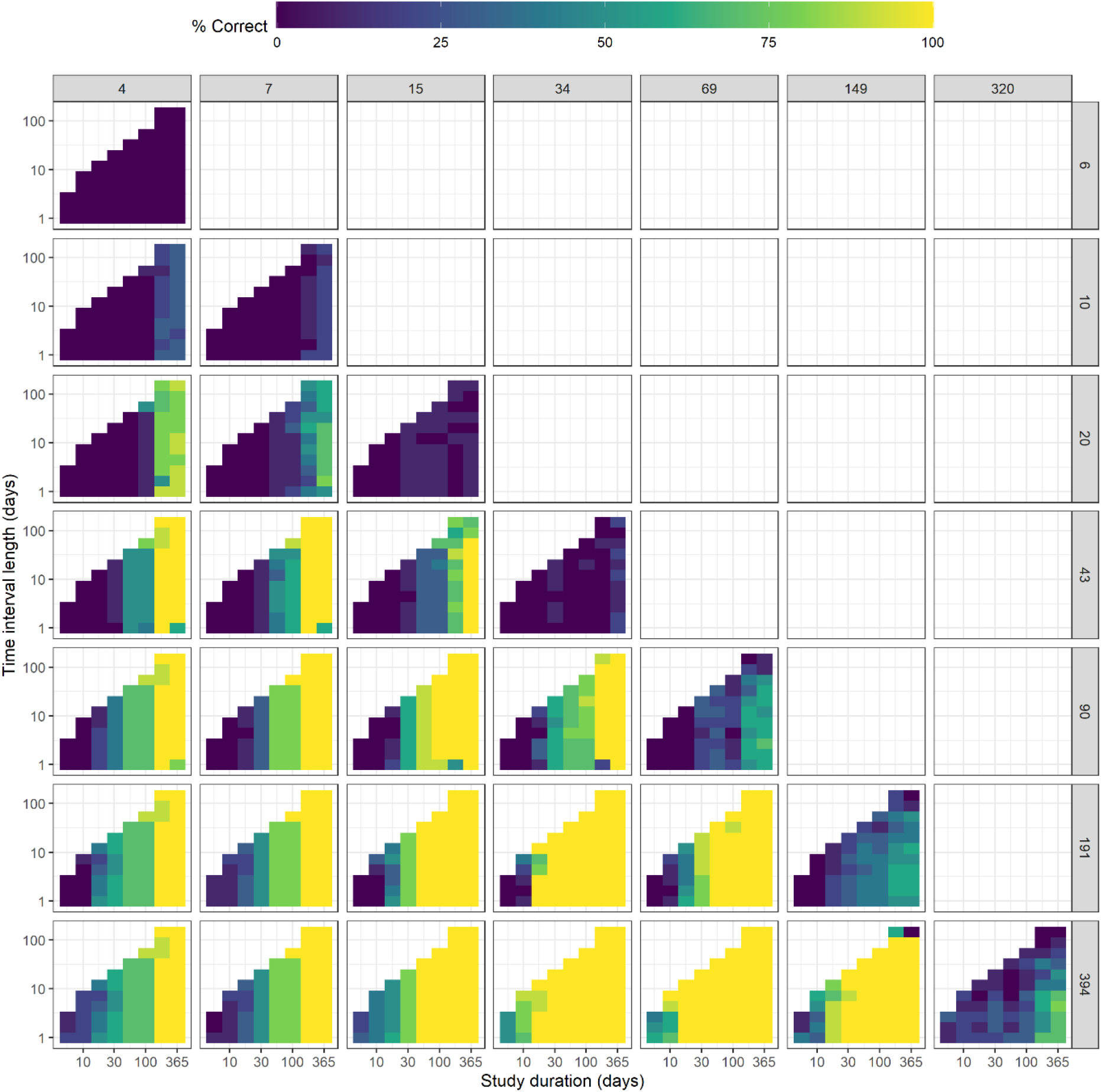
The percentage of times the RN model resulted in a significant positive effect of location on estimated abundance (where location B (rows) has a higher abundance than location A (columns)), per sampling period (days) and interval length (days). In this figure, only results with 25 cameras per location are shown.

The z-values of the RN models were estimated in the multivariate polynomial regression (using the total sampling effort (i.e. the cumulative number of camera trapping days of all used camera traps), the proportion of camera traps with detections, the number of camera traps, and the interval length) with an adjusted R^2^ of 0.583 (Table S1). The z-value is (non-linearly) positively related with both sampling effort and *P*_*presence*_ (Fig. 2A). In case of small sampling effort and/or small proportions of cameras with detections, the z-value is below 1.96, indicating that the RN model cannot compute reliable abundance estimates, regardless of the interval length used.

**Figure 2:**
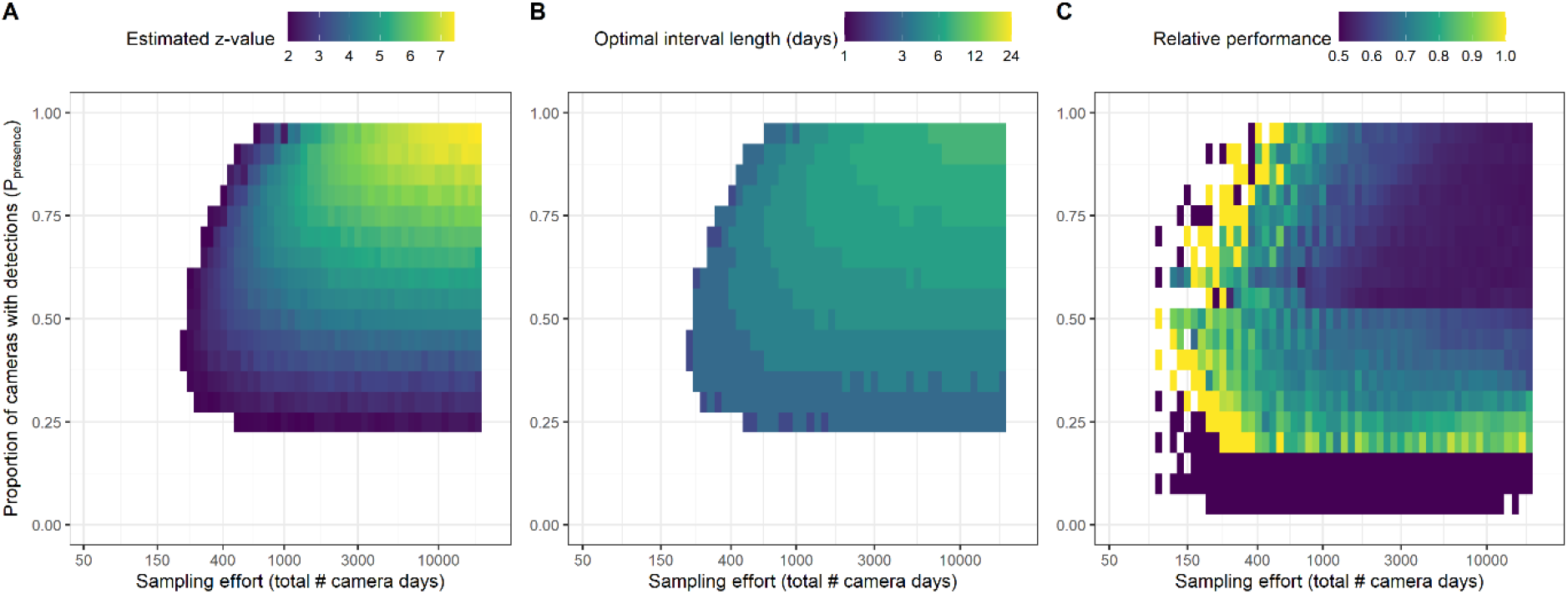
The estimated z-value of the best-fitting Royle-Nichols occupancy model (A), the corresponding interval length (in days) that was used to obtain this optimal fit (B), and the relative performance of the best-fitting model over the model that uses a 5-day interval (C), per sampling effort (i.e. the total number of camera days) and proportion of cameras with detection. The relative performance is calculated as p_opt_ / (p_5_ + p_opt_), where p_opt_ is the percentage correct at the optimal time interval and p_5_ is the percentage of correct results when using a 5-day interval. In (A) and (B), only results with z > 1.96 are shown.

The optimal interval length that corresponds to the highest z-values per sampling effort and proportion of cameras with detections also relates to these parameters. Large interval lengths (> 10 days) should be used in case of high sampling effort (> 1,000 days) and in case of a small proportion of cameras with detections of the species (*P*_*presence*_ < 0.5; Fig. 2B). In contrast, the interval length should be small (< 3 days) when *P*_*presence*_ is high to avoid that the matrix consists almost exclusively of ones, or when sampling effort is small.

We found that the RN model that uses the optimal interval length does not always provide better results than a model with a 5-day interval (Fig. 2C). When *P*_*presence*_ and/or sampling effort are low, none of the RN models results in correct conclusions. On the other hand, when both *P*_*presence*_ and sampling effort are high, RN models almost always result in correct answers, regardless of the interval length used. However, for the entire parameter space in between these two extremes, using the optimized interval length substantially improved the fit of the RN model to the data (Fig. 2C).

## 3. Optimizing segmentation

Given the above-mentioned results, guidelines can be generated for choosing the optimum segmentation for comparing densities using RN models, depending on animal density and study design. Here, we developed a tool to do so. We fitted a polynomial model (Supplement A) to estimate the z-value from data that can easily be obtained from camera trapping records: (i) sampling effort (*N*_*SE*_), (ii) proportion of cameras with detections (*P*_*presence*_), (iii) number of cameras (*N*_*cams*_), and (iv) interval length (*dT*). As field data often contains different recording durations per camera trap, we used total sampling effort (*N*_*SE*_, i.e. the total number of camera days) instead of the parameter study duration in this polynomial model. To ensure that the z-value that results from extrapolated, high sampling efforts will not become excessively large, we fit a linear (negative) relation between the inverse of the sampling effort and the z-value. A second parameter in the polynomial model is the proportion of cameras with detections (*P*_*presence*_). In most field studies, the actual density per location and per species is unknown. As the proportion of cameras with detections relates well with average density and study duration in our model simulations (Fig. S2), we used the logit transformation of this parameter (as *P*_*presence*_ ranges between 0 and 1) as a proxy of average abundance. The third and fourth parameter we used in the polynomial model are the number of cameras used in the study (*N*_*cams*_) and the interval length (*dT*) used to assemble the data for analysis with the RN model. We assumed a linear (negative) relation between the inverse of the number of cameras used and the z-value.

Subsequently, we generated an R-function that uses the above estimation of the z-value to calculate the optimal interval length (*dT*) given camera-trap data on a certain species (Fig. S3; De Jager 2024). The input is a simple presence/absence matrix, the output is the estimated optimal interval length and the minimum number of intervals needed per camera (i.e. number of columns in the matrix). The presence/absence matrix consists of records per day (columns) and camera trap (rows), with three possible values: one (at least one detection that day at that camera trap), zero (no detection), or NA (sampling has stopped). The fit of the RN model is then calculated for a large range of combinations of the minimum number of intervals needed per camera (2 ≤ *nT* ≤ 30) and interval lengths (1 ≤ *dT* < *dT*_*max*_ ; where *dT*_*max*_ is the 95 percentile of survey effort per camera divided by *nT*). For each combination of *nT* and *dT*, a new presence/absence matrix is made, with which the total survey effort, number of cameras used, and the proportion of cameras with detections is calculated. These values of *N*_*SE*_, *N*_*cams*_, and *P*_*presence*_, together with *dT*, are then used to estimate the fit of the RN model, where the fit equals the inverse of the estimated z-value (1/e^z^). Finally, the values of *dT* and *nT* are selected that correspond to the fit-value closest to zero.

## 4. Field application: Forest mammals in the Republic of Congo and Gabon

We evaluated how segmentation optimization affected the power of occupancy-based site comparisons by applying it to camera trapping data from Zwerts et al. (2024), collected in tropical forests across the Republic of Congo and Gabon. In this study, seven pairs of non-certified and FSC-certified timber logging concessions (timber logged according to principles of the Forest Stewardship Council, FSC) were surveyed with camera traps during 2018 – 2021, using 474 camera-trap locations in total to examine whether FSC-certification benefits animals in logged forests. In Zwerts *et al*. (2024), relative abundance was analyzed using encounter rates.

For each of the 21 mammalian species with sufficient data (> 150 detections), we created a presence/absence matrix that contained daily detections per camera, starting with the first camera trapping day in the first column. Row length depended on the survey effort (in days) of the camera with the highest number of camera trapping days, and column length depended on the number of cameras used in the study. Each cell (*i*,*j*) of the matrix contained the values 1 for detection of the species by camera *i* on day *j*, 0 for non-detection of the species by camera *i* on day *j*, and NA in case the camera had already stopped recording at that day.

First, we estimated the optimal interval length (*dT*) and the minimum number of intervals that were needed per camera (*nT*) for each species. These parameters determine how many cameras are to be included (*N*_*cams*_, cameras with too little sampling effort should be disregarded), and hence also affect the total sampling effort (*N*_*SE*_) and the proportion of cameras with detections (*P*_*presence*_). We calculated *N*_*cams*_, *N*_*SE*_, and *P*_*presence*_ for all possible combinations of *dT* and *nT*. Using the polynomial model (Suppl. A, eq. S1), we estimated how well the RN model should be able to provide correct results; the interval length and minimum number of intervals needed per camera that corresponded to the lowest fit-value were used in the further analysis of the data.

The RN model estimated the difference in relative abundance of a species between forest management types. For the field data analysis, detection probability at each camera trapping location *i* was assumed to be positively affected by visibility (*v*; which was categorized into 0-10 m, 11-20 m, and > 20 m):

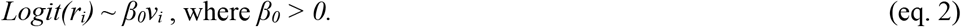

The abundance estimate (*λ*_*i*_) of a species was calculated per location-pair (*Loc* a to g) and area type (*A*: FSC versus non-FSC):

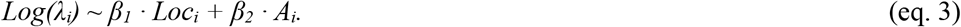

The RN model provided, amongst others, a z-statistic for the probability that these abundances are equal at FSC-certified and non-FSC concessions. When z > 1.96, a species’ abundance was estimated to be significantly higher in FSC-certified than in non-FSC concessions (p < 0.05); likewise, when z < -1.96, a species’ abundance was estimated to be significantly lower in FSC-certified than in non-FSC concessions.

As is often the case in camera-trapping studies, the sampling period per camera varied widely, ranging between 1 and 239 days per camera. Depending on the interval length and the minimum number of intervals required per camera, the total sampling effort thus differed greatly, as all cameras with a shorter individual sampling effort than *dT* x *nT* were disregarded (Fig. S4). Consequently, the proportion of cameras with detections of a species varied with interval length and minimum sample size as well (Fig. 3 top panels).

**Figure 3:**
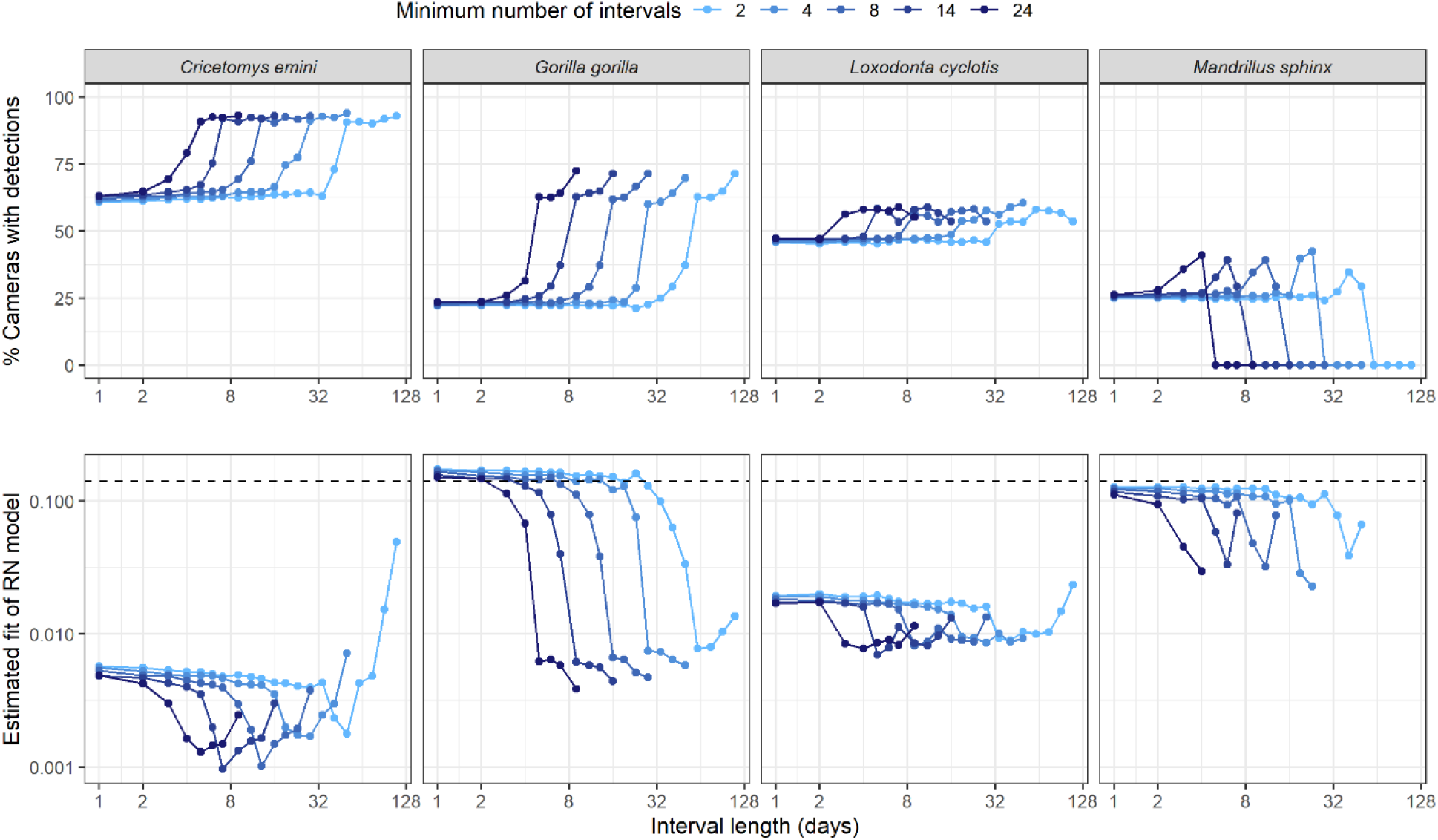
Dependence of animal detection on interval length and number for four mammals in the forests of the Republic of Congo and Gabon. Graphs show the proportion of cameras with detections (top panels) and the estimated fit of the Royle-Nichols occupancy model (bottom panels) as a function of interval length (in days) and minimum sample size (colors). The dashed lines in the bottom panels indicate the threshold value below which the RN model is estimated to provide adequate results (closest to zero = best fit).

Using the optimization method, we selected the segmentation and minimum number of intervals per species that corresponded to the estimated fit-values of the RN model (= 1/e^z^) that were closest to zero (Fig. 3 bottom panels). The resulting interval lengths and minimum sample sizes varied substantially between species (Fig. 4). Optimal interval lengths ranged between 3 and 70 days, while the minimum number of intervals ranged between 2 and 30, which comes down to 10 to 230 sampling days.

**Figure 4:**
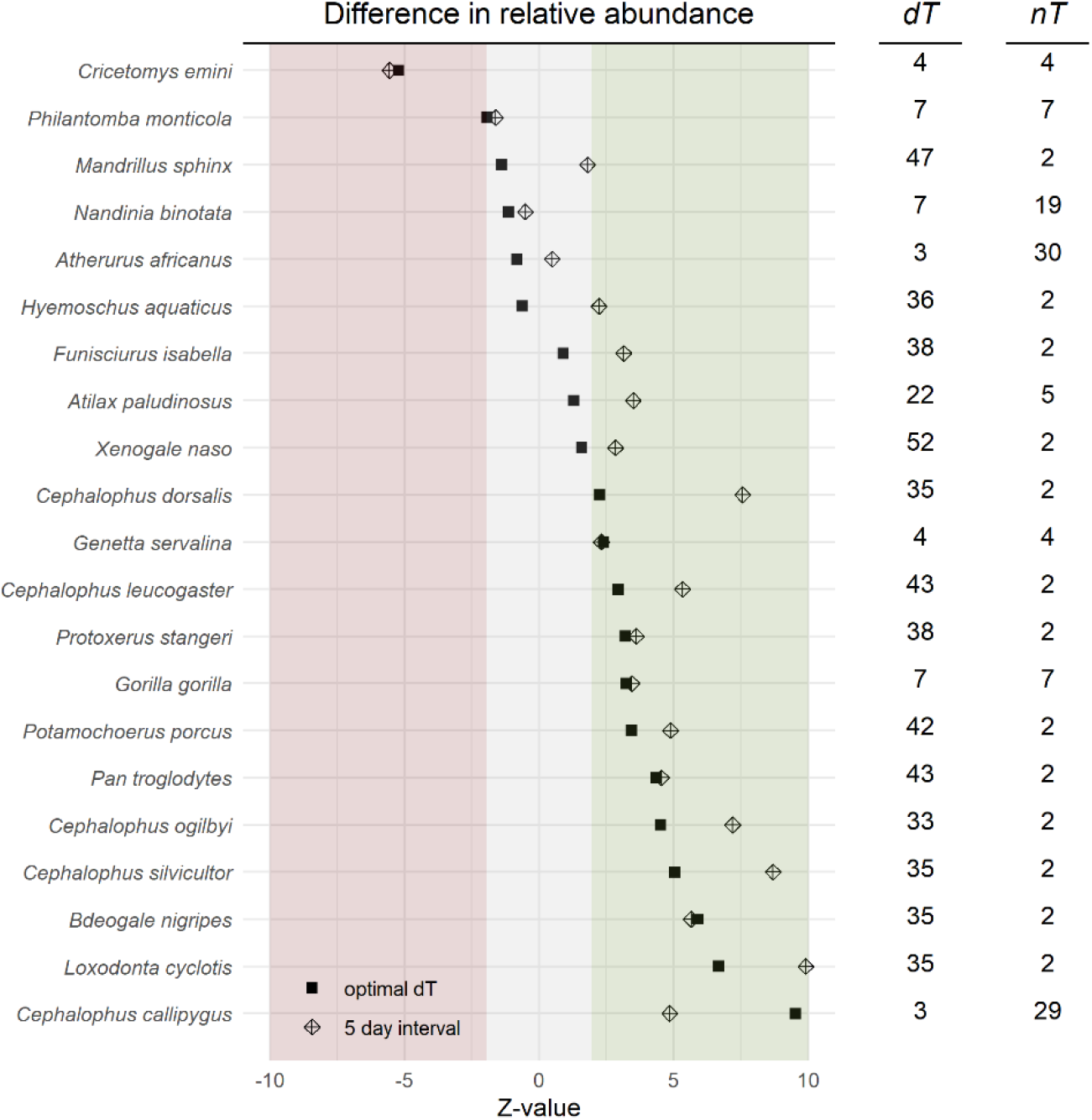
Comparison of occupancy between concessions in the Republic of Congo and Gabon with and without FSC certification, for 21 mammal species. Z-values, resulting from the Royle-Nichols abundance model using the optimal interval length (squares) or the commonly used 5-day interval (diamonds), indicating whether a species is estimated to be more abundant in FSC-certified forests than in non-FSC concessions (green area, z > 1.96), more abundant in non-FSC than in FSC-certified concessions (red area, z < -1.96), or equally abundant in both FSC and non-FSC concessions (grey area, -1.96 ≤ z ≤ 1.96). The columns show the optimal time interval length (dT, in days) and the corresponding minimum number of intervals (nT).

Comparing these results to those produced with the arbitrarily chosen standard 5-day interval shows that the models with optimized interval lengths resulted in fewer significant outcomes (Fig. 4). That is, for most species, the z-value estimated with the optimized RN model was closer to zero (e.g. more precautious) than those estimated using a 5-day interval. With the 5-day interval, an additional 5 species were estimated to be more abundant in FSC-certified concessions than in non-FSC concessions. For 4 rather than 8 species, there was no significant difference. The model with optimized intervals thus yielded more conservative results.

Results of the RN model varied greatly between species and were significantly affected by the choice of interval length and minimum number of intervals (Fig. 5; Fig. S5). In most cases, the z-value that estimated the difference in relative abundance between FSC-certified and non-FSC concessions was less significant as the interval was longer. For some species, a significant positive effect of FSC-certified management can be found on relative abundance when using certain interval lengths, while this can switch to a significant negative effect when the interval length is altered.

**Figure 5:**
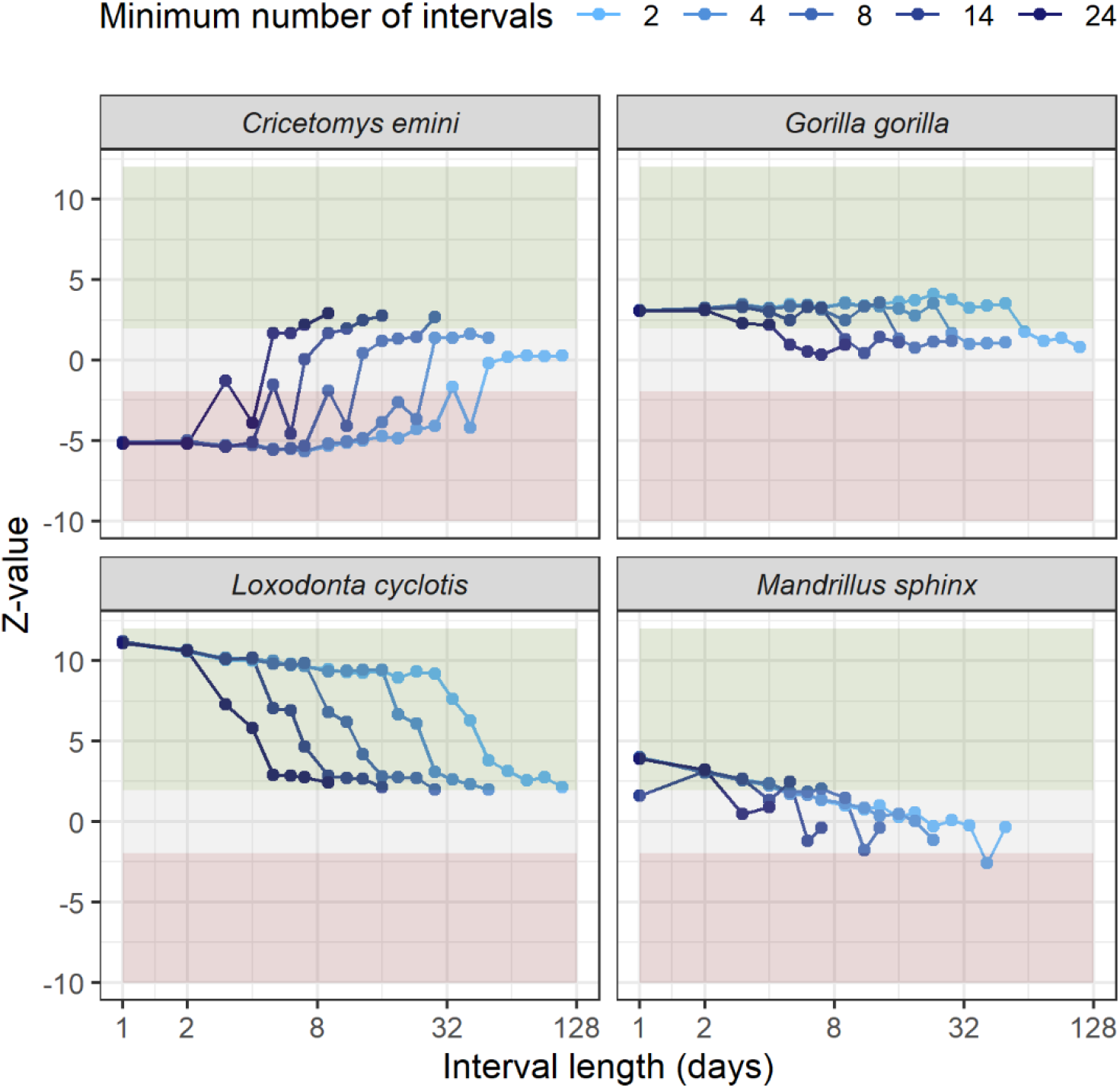
Dependence of model outcomes on the length and number of intervals for four species of mammal in the rainforests of the Republic of Congo and Gabon. Plotted are the Z-values resulting from RN models for different interval lengths (in days) and minimum sample sizes (colors). Z-values indicate whether a species is estimated to be more abundant in FSC-certified concessions than in non-FSC concessions (green area, z > 1.96), in non-FSC than in FSC-certified concessions (red area, z < -1.96), or equally abundant in both concession types (grey area, -1.96 ≤ z ≤ 1.96).

## 5. Discussion

Segmentation, i.e., choosing the number and length of sampling intervals, is a necessary step in applying the Royle-Nichols occupancy model to camera-trapping data, but guidelines for optimizing this choice have been lacking. Here, we determined how segmentation affected the ability of the RN model to estimate whether relative abundances significantly differ from one another, using camera trapping events of simulated populations with known densities. We found that segmentation can have a substantial impact on the accuracy of the RN model’s results. Using these findings, we developed guidelines for optimizing segmentation.

Application of the optimization to camera-trapping data from a field study that compared wildlife between FSC-certified and non-FSC forests in the Republic of Congo and Gabon (Zwerts et al. 2024) revealed that the choice of interval length can affect the direction of relationships. For some species in this study, we found significant negative effects of FSC-management on relative abundance when using certain interval lengths, and significant positive effects when using other interval lengths. This indicates that the interval length is by no means a parameter to be standardized at a given value but should be carefully chosen based on the properties of the data at hand.

We found that study duration positively affected the capability of the RN model to detect differences in relative abundance between populations (Fig. 1). An exception here is when densities are too high and hence the number of detections too large (excess of ‘ones’ in the data) for the RN model to handle, but this could be remedied by trimming the dataset. Likewise, the number of cameras used in a study positively affected the success of the RN model. Thus, the RN model is more likely to correctly estimate differences in relative abundance between locations as the studies are longer and involve more sampling points.

MacKenzie and Royle (2005) suggested that, for rare species, it is more efficient to survey more sampling units less intensively, while for common species, fewer sampling units should be surveyed more intensively. In general, logically, our predictions of the RN model were more accurate for longer studies, for higher average densities, and for larger differences in densities between study locations. In practice, however, study length and number of sampling points are constrained by the availability of equipment, labour and battery power (Zwerts et al. 2021); other studies have provided guidelines and tools to optimize that choice (e.g., Beaudrot et al. 2019, Kays et al. 2020, Zwerts et al. 2021).

In camera trapping, a given study duration can be segmented into as many sampling units as it contains days, and many studies choose this option as to retain as much information as possible. We however find that there is an optimum interval number and length for detecting differences in occupancy. In short studies (< 30 days), six or more short time intervals (< 5 days) yielded better estimates than fewer long time intervals (if average densities are sufficiently high), hence the choice for having intervals of one day works well. In longer studies, however, high average wildlife densities were best predicted with a large number of short intervals (< 5 days) to avoid occupancy values of 1 at all sites, but low average densities were best predicted with fewer, long intervals (>> 5 days). Thus, an interval length of one day is not always the best choice. The optimal interval length and number is thus species specific, and dependent on rarity. Our study provides a tool for optimization of interval lengths.

Our study has a number of possible limitations. The most important one is that our assessment of the effects of segmentation was based on simulated data that do not reflect the complexity of animal movement. In particular, in most species, individuals constrain their movements to ranges that are much smaller than the survey area. This may lead to concentrations of detections at a subset of the sampling points, with some points detecting many animals and others few or none. With increasingly skewed numbers of detections across camera trap locations, the need to replicate over space rather than time increases. Constrained movements of individuals would obviously change the patterns that we found, but the need for optimization remains the same. It should also be noted that techniques of occupancy modelling are still advancing rapidly. One particularly noteworthy development is the idea of dropping the subdivision of camera-trap data altogether, and instead to treat the detection process as occurring in continuous time. This does not only omit the need for choosing the proper subdivision, but also allows studying species interactions at a much finer scale (Kellner et al. 2022). Kellner et al.(2022) recently developed an occupancy model for continuous time and applied it to camera-trap data (see also Parsons et al. 2022).

This study unequivocally demonstrated that segmentation , i.e., the choice of interval size and number, plays a pivotal role in the accuracy of results in occupancy modelling. The impact of this decision cannot be overstated, as it has far-reaching implications for ecological research and the assessment of conservation efforts. However, there is no one-size-fits-all solution. The optimal segmentation is inherently context-dependent, and may depend on a myriad of factors such as the species’ behavior, the spatial and temporal scales of the study and the intricacies of the modelling techniques employed. We therefore advocate for a meticulous approach for segmentation, coupled with the implementation of a well-designed sampling strategy.

For the future, we suggest to develop systematic frameworks for data segmentation that can be adapted across diverse ecological contexts. Investigating how different species, habitats, and environmental conditions interact with segmentation will be instrumental in establishing generalizable guidelines. Furthermore, to enhance efficiency in the selection process, future studies could explore the integration of advanced statistical techniques and machine learning algorithms to assist in automating the selection process. Recognizing the broader implications, our recommendation extends to the evaluation of conservation efforts. The effectiveness of conservation initiatives hinges on the accurate assessment of species abundance, making the careful selection of interval size a linchpin in the evaluation of such initiatives. In conclusion, our study underscores the significance of thoughtful segmentation of camera-trapping data for occupancy modelling.

## Acknowledgements

MJ, MK, and PJ received funding by the Institutional Collaboration Grant from the graduate school Production Ecology and Resource Conservation (PE&RC).

## Tables

**Table 1:**
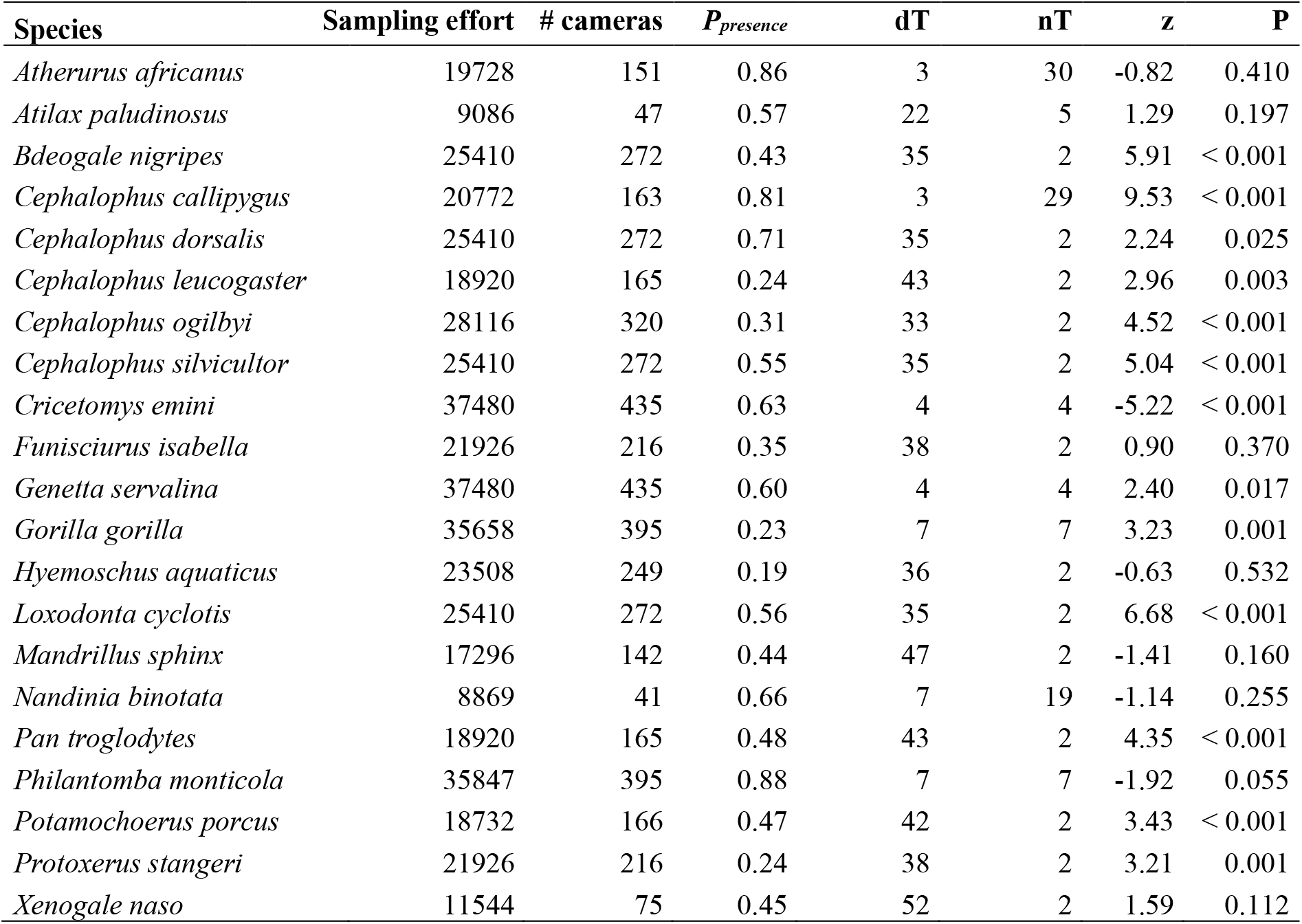
Per species, we estimated the optimal interval length for the RN occupancy analysis between FSC and non-FSC forests. The corresponding sampling effort (i.e. total number of camera days), the number of cameras used, the proportion of cameras with detections (P_presence_), the optimal interval length (dT, in days), the minimum number of intervals allowed per camera (nT), the z-value (values > 1.96 indicate higher abundance in FSC than in non-FSC forests), and significance (P) are shown.

| Species | Sampling effort | # cameras | $P_{\text{presence}}$ | dT | nT | z | P |
| --- | --- | --- | --- | --- | --- | --- | --- |
| *Atherurus africanus* | 19728 | 151 | 0.86 | 3 | 30 | -0.82 | 0.410 |
| *Atilax paludinosus* | 9086 | 47 | 0.57 | 22 | 5 | 1.29 | 0.197 |
| *Bdeogale nigripes* | 25410 | 272 | 0.43 | 35 | 2 | 5.91 | < 0.001 |
| *Cephalophus callipygus* | 20772 | 163 | 0.81 | 3 | 29 | 9.53 | < 0.001 |
| *Cephalophus dorsalis* | 25410 | 272 | 0.71 | 35 | 2 | 2.24 | 0.025 |
| *Cephalophus leucogaster* | 18920 | 165 | 0.24 | 43 | 2 | 2.96 | 0.003 |
| *Cephalophus ogilbyi* | 28116 | 320 | 0.31 | 33 | 2 | 4.52 | < 0.001 |
| *Cephalophus silvicultor* | 25410 | 272 | 0.55 | 35 | 2 | 5.04 | < 0.001 |
| *Cricetomys emini* | 37480 | 435 | 0.63 | 4 | 4 | -5.22 | < 0.001 |
| *Funisciurus isabella* | 21926 | 216 | 0.35 | 38 | 2 | 0.90 | 0.370 |
| *Genetta servalina* | 37480 | 435 | 0.60 | 4 | 4 | 2.40 | 0.017 |
| *Gorilla gorilla* | 35658 | 395 | 0.23 | 7 | 7 | 3.23 | 0.001 |
| *Hyemoschus aquaticus* | 23508 | 249 | 0.19 | 36 | 2 | -0.63 | 0.532 |
| *Loxodonta cyclotis* | 25410 | 272 | 0.56 | 35 | 2 | 6.68 | < 0.001 |
| *Mandrillus sphinx* | 17296 | 142 | 0.44 | 47 | 2 | -1.41 | 0.160 |
| *Nandinia binotata* | 8869 | 41 | 0.66 | 7 | 19 | -1.14 | 0.255 |
| *Pan troglodytes* | 18920 | 165 | 0.48 | 43 | 2 | 4.35 | < 0.001 |
| *Philantomba monticola* | 35847 | 395 | 0.88 | 7 | 7 | -1.92 | 0.055 |
| *Potamochoerus porcus* | 18732 | 166 | 0.47 | 42 | 2 | 3.43 | < 0.001 |
| *Protoxerus stangeri* | 21926 | 216 | 0.24 | 38 | 2 | 3.21 | 0.001 |
| *Xenogale naso* | 11544 | 75 | 0.45 | 52 | 2 | 1.59 | 0.112 |

