## Supplementary Information for "OPTIMIZING SEGMENTATION IN OCCUPANCY MODELLING OF CAMERA-TRAP DATA"

**Contents:** **page number**

| | |
|---|---|
| Supplement A: polynomial model | 2 |
| Supplementary Table S1 | 3 |
| Supplementary Figure S1 | 4 |
| Supplementary Figure S2 | 5 |
| Supplementary Figure S3 | 6 |
| Supplementary Figure S4 | 7 |
| Supplementary Figure S5 | 8 |

**Supplement A: Polynomial model**

We fitted a polynomial model to estimate the z-value from data that can easily be obtained from camera trapping observations: (i) sampling effort ($N_{SE}$), (ii) proportion of cameras with detections ($P_{presence}$), (iii) number of cameras ($N_{cams}$), and (iv) interval length ($dT$). As field data often contains different recording durations per camera trap, we used total sampling effort ($N_{SE}$, i.e. the total number of camera days) instead of the parameter study duration in this polynomial model. To ensure that the z-value that results from extrapolated, high sampling efforts will not become excessively large, we fit a linear (negative) relation between the inverse of the sampling effort and the z-value. A second parameter in the polynomial model is the proportion of cameras with detections ($P_{presence}$). In most field studies, the actual density per location and per species is unknown. As the proportion of cameras with detections relates well with average density and study duration in our model simulations (Fig. S1), we used the logit transformation of this parameter (as $P_{presence}$ ranges between 0 and 1) as a proxy of average abundance. The third and fourth parameter we used in the polynomial model are the number of cameras used in the study ($N_{cams}$) and the interval length ($dT$) used to assemble the data for analysis with the RN model. We assumed a linear (negative) relation between the inverse of the number of cameras used and the z-value. The z-value was thus calculated as:

$$\begin{aligned}
z \sim\ & \beta_1 + \beta_2\, dT + \beta_3\, N_{SE}^{-1} + \beta_4\, Logit(P_{presence}) + \beta_5\, N_{cams}^{-1} + \\
& \beta_6\, dT^2 + \beta_7\, Logit(P_{presence})^2 + \beta_8\, dT^3 + \beta_9\, Logit(P_{presence})^3 + \\
& \beta_{10}\, dT \cdot N_{SE}^{-1} + \beta_{11}\, dT \cdot Logit(P_{presence}) + \beta_{12}\, N_{SE}^{-1} \cdot Logit(P_{presence}) + \beta_{13}\, dT \cdot N_{cams}^{-1} + \\
& \beta_{14}\, N_{SE}^{-1} \cdot N_{cams}^{-1} + \beta_{15}\, Logit(P_{presence}) \cdot N_{cams}^{-1} + \beta_{16}\, N_{SE}^{-1} \cdot dT^2 + \beta_{17}\, dT^2 \cdot Logit(P_{presence})^2 + \\
& \beta_{18}\, N_{SE}^{-1} \cdot Logit(P_{presence})^2 + \beta_{19}\, N_{cams}^{-1} \cdot dT^2 + \beta_{20}\, N_{cams}^{-1} \cdot Logit(P_{presence})^2 + \beta_{21}\, N_{SE}^{-1} \cdot dT^3 + \\
& \beta_{22}\, dT^3 \cdot Logit(P_{presence})^3 + \beta_{23}\, N_{SE}^{-1} \cdot Logit(P_{presence})^3 + \beta_{24}\, N_{cams}^{-1} \cdot dT^3 + \\
& \beta_{25}\, N_{cams}^{-1} \cdot Logit(P_{presence})^3 + \beta_{26}\, dT \cdot N_{SE}^{-1} \cdot Logit(P_{presence}) + \beta_{27}\, dT \cdot N_{SE}^{-1} \cdot N_{cams}^{-1} + \\
& \beta_{28}\, dT \cdot Logit(P_{presence}) \cdot N_{cams}^{-1} + \beta_{29}\, N_{SE}^{-1} \cdot Logit(P_{presence}) \cdot N_{cams}^{-1} + \\
& \beta_{30}\, N_{SE}^{-1} \cdot dT^2 \cdot Logit(P_{presence})^2 + \beta_{31}\, N_{SE}^{-1} \cdot N_{cams}^{-1} \cdot dT^2 + \beta_{32}\, N_{cams}^{-1} \cdot dT^2 \cdot Logit(P_{presence})^2 + \\
& \beta_{33}\, N_{SE}^{-1} \cdot N_{cams}^{-1} \cdot Logit(P_{presence})^2 + \beta_{34}\, N_{SE}^{-1} \cdot dT^3 \cdot Logit(P_{presence})^3 + \beta_{35}\, N_{SE}^{-1} \cdot N_{cams}^{-1} \cdot dT^3 + \\
& \beta_{36}\, N_{cams}^{-1} \cdot dT^3 \cdot Logit(P_{presence})^3 + \beta_{37}\, N_{SE}^{-1} \cdot N_{cams}^{-1} \cdot Logit(P_{presence})^3 + \\
& \beta_{38}\, dT \cdot N_{SE}^{-1} \cdot Logit(P_{presence}) \cdot N_{cams}^{-1} + \beta_{39}\, N_{SE}^{-1} \cdot N_{cams}^{-1} \cdot dT^2 \cdot Logit(P_{presence})^2 + \\
& \beta_{40}\, N_{SE}^{-1} \cdot N_{cams}^{-1} \cdot dT^3 \cdot Logit(P_{presence})^3 \qquad \text{(eq. A1)}
\end{aligned}$$

**Table S1:** Coefficients (see Eq. 1), coefficient estimates, standard errors (SE), t-values, and P-values of the multivariate polynomial regression model that estimates the z-values of the RN models from the total sampling effort, the proportion of camera traps with detections, the number of camera traps, and the interval length.

| **Coefficient** | **Estimate** | **SE** | **t-value** | **P-value** |
|---|---:|---:|---:|---:|
| β1 | 4.54E+00 | 2.31E-03 | 1966.45 | < 0.001 |
| β2 | 7.92E-03 | 1.98E-04 | 39.99 | < 0.001 |
| β3 | -1.65E+03 | 3.50E+00 | -470.72 | < 0.001 |
| β4 | 2.11E+00 | 1.70E-03 | 1239.94 | < 0.001 |
| β5 | -3.61E+01 | 5.20E-02 | -693.33 | < 0.001 |
| β6 | 4.44E-05 | 3.87E-06 | 11.47 | < 0.001 |
| β7 | -8.92E-02 | 4.64E-04 | -192.18 | < 0.001 |
| β8 | -3.94E-07 | 1.94E-08 | -20.26 | < 0.001 |
| β9 | -6.40E-02 | 2.12E-04 | -301.24 | < 0.001 |
| β10 | -3.57E+01 | 5.02E-01 | -71.15 | < 0.001 |
| β11 | -2.50E-03 | 4.55E-05 | -55.05 | < 0.001 |
| β12 | -1.99E+03 | 2.88E+00 | -692.36 | < 0.001 |
| β13 | -1.08E-01 | 4.52E-03 | -23.95 | < 0.001 |
| β14 | 1.71E+04 | 4.85E+01 | 352.77 | < 0.001 |
| β15 | -5.84E+00 | 4.14E-02 | -140.89 | < 0.001 |
| β16 | -6.99E-01 | 1.76E-02 | -39.70 | < 0.001 |
| β17 | -3.59E-05 | 3.20E-07 | -111.96 | < 0.001 |
| β18 | -7.00E+02 | 1.29E+00 | -544.20 | < 0.001 |
| β19 | -6.11E-04 | 8.72E-05 | -7.01 | < 0.001 |
| β20 | 1.42E+00 | 1.24E-02 | 114.22 | < 0.001 |
| β21 | 2.76E-03 | 1.39E-04 | 19.85 | < 0.001 |
| β22 | 4.91E-08 | 7.79E-10 | 63.03 | < 0.001 |
| β23 | -7.77E+01 | 2.90E-01 | -267.80 | < 0.001 |
| β24 | 7.49E-06 | 4.55E-07 | 16.45 | < 0.001 |
| β25 | -1.05E+00 | 6.46E-03 | -162.45 | < 0.001 |
| β26 | -1.43E+01 | 1.82E-01 | -78.30 | < 0.001 |
| β27 | 3.55E+02 | 6.97E+00 | 50.96 | < 0.001 |
| β28 | 4.38E-02 | 1.11E-03 | 39.53 | < 0.001 |
| β29 | 1.94E+04 | 4.14E+01 | 468.66 | < 0.001 |
| β30 | 4.60E-02 | 1.84E-03 | 25.03 | < 0.001 |
| β31 | 1.06E+01 | 2.32E-01 | 45.43 | < 0.001 |
| β32 | 6.05E-04 | 7.78E-06 | 77.84 | < 0.001 |
| β33 | 6.96E+03 | 1.91E+01 | 364.30 | < 0.001 |
| β34 | -4.39E-05 | 6.57E-06 | -6.68 | < 0.001 |
| β35 | -5.34E-02 | 1.58E-03 | -33.81 | < 0.001 |
| β36 | -7.25E-07 | 2.42E-08 | -29.99 | < 0.001 |
| β37 | 9.41E+02 | 5.51E+00 | 170.80 | < 0.001 |
| β38 | 7.90E+01 | 2.61E+00 | 30.29 | < 0.001 |
| β39 | -1.19E+00 | 2.71E-02 | -43.97 | < 0.001 |
| β40 | 7.20E-05 | 9.38E-05 | 0.77 | 0.442441 |

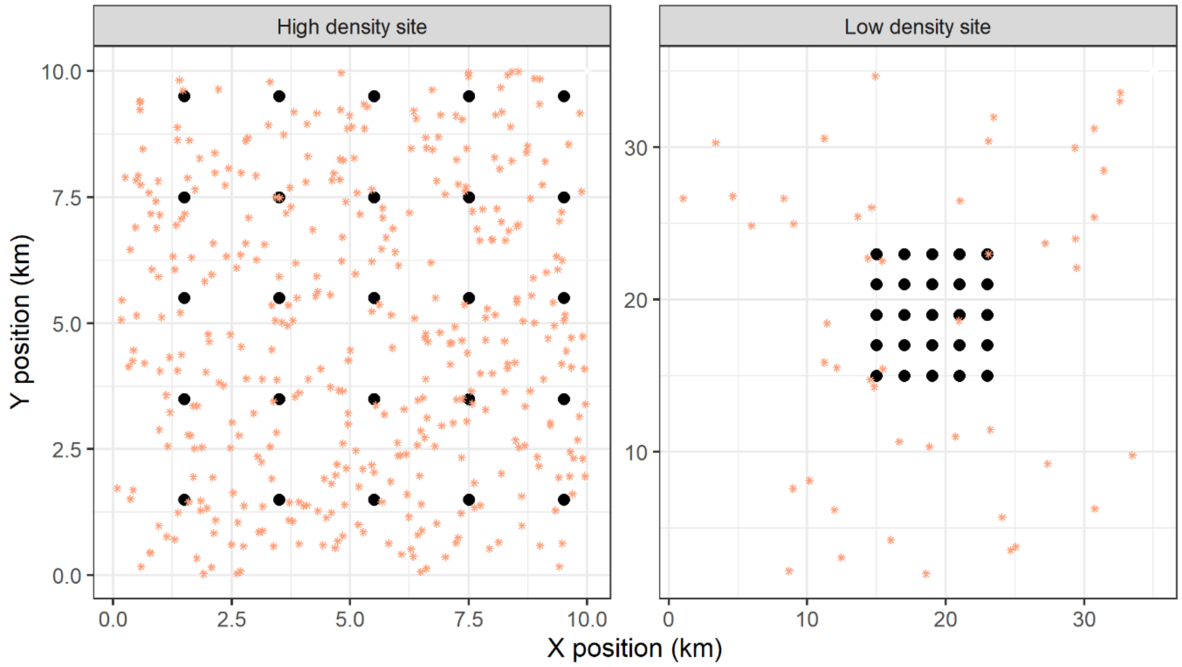


***Figure S1:*** *Illustration of two scenarios in the individual-based simulation model. The black squares represent 25 camera traps that detect individuals passing through 5 x 5 m areas within a 10 x 10 km square. The asterisks represent individual animals that are randomly placed in the simulated area (grey) and move around. The high and low density scenario contain 4 and 0.04 individuals km$^{-2}$, respectively. As the boundaries are periodic, the location of the camera grid within the simulation area is irrelevant.*

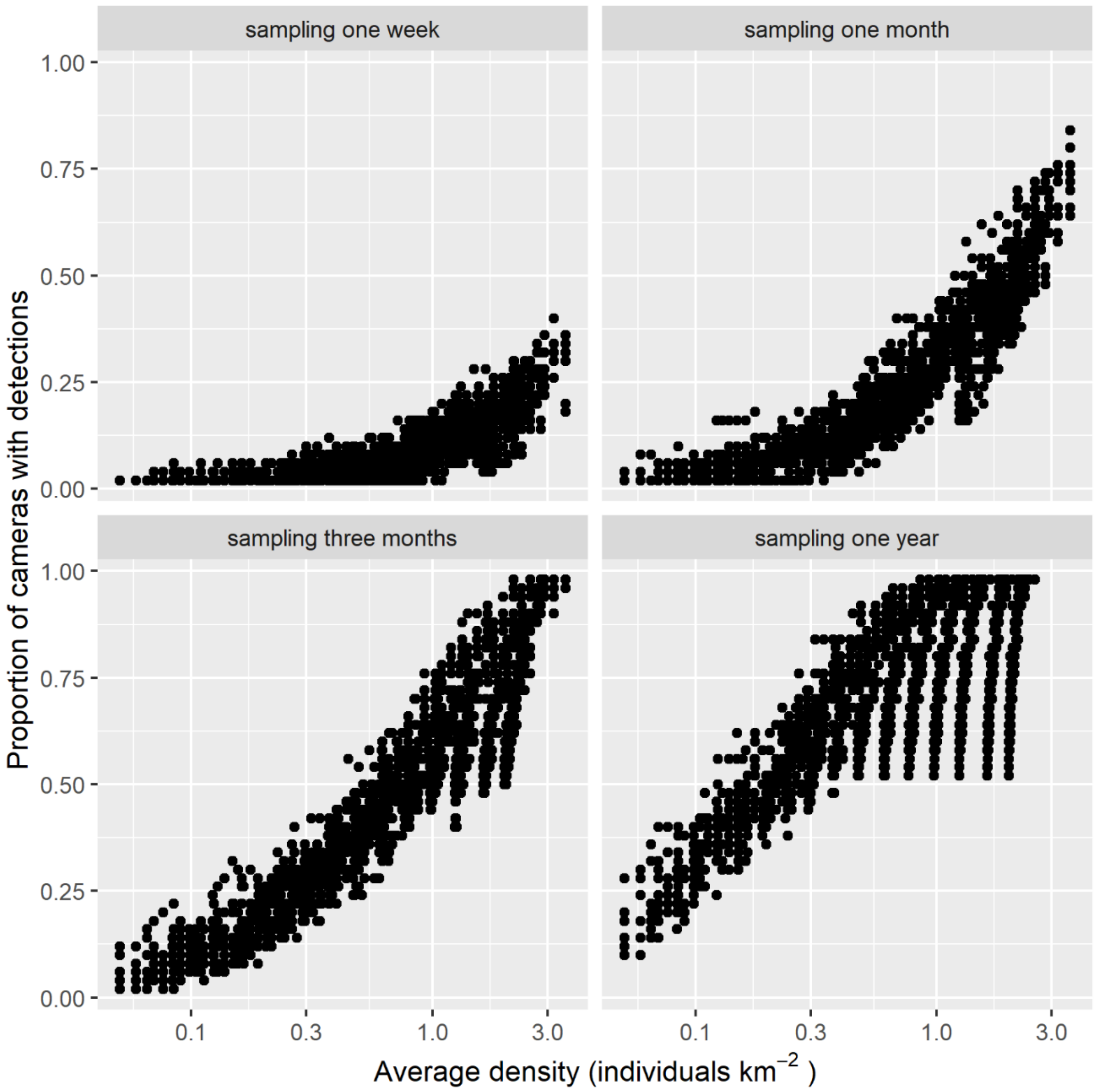


***Figure S2:*** *The proportion of cameras with detections ($P_{presence}$) relates well with the average density of individuals at the two simulated locations. The relation changes with sampling period, as more cameras are able to detect the species given more sampling time.*

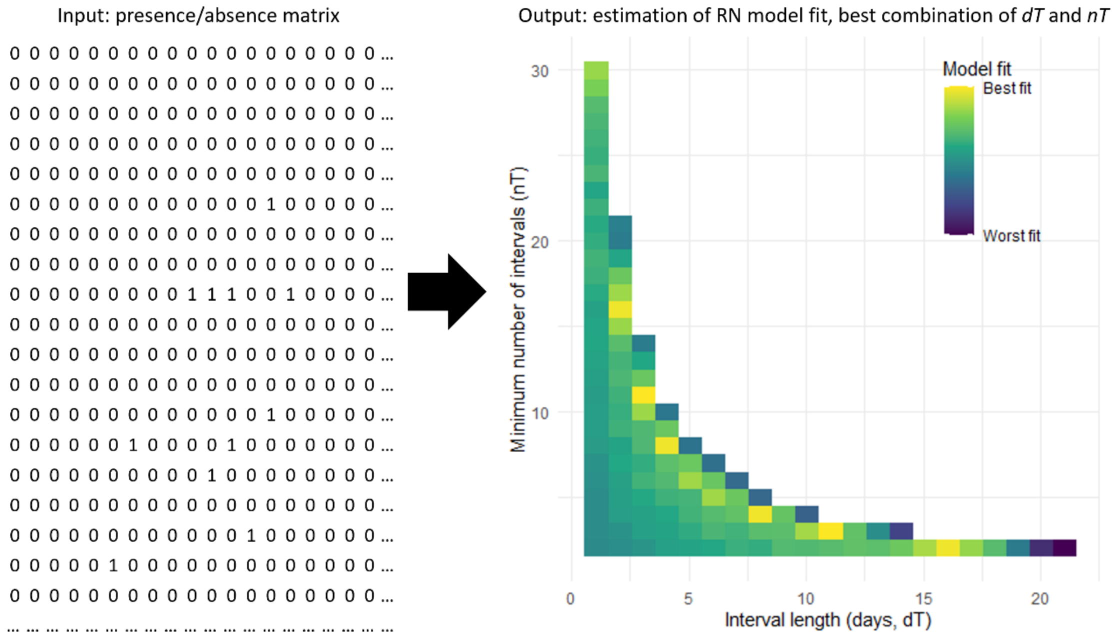


***Figure S3:*** *Illustration of how the interval length (dT) and minimum number of intervals (nT) are estimated from presence/absence input data. Using the polynomial regression function (Suppl. A; Table S1;* *https://doi.org/10.5281/zenodo.10424189* *), the likelihood that the RN model will provide accurate estimations of differences in relative abundances between locations is estimated in the form of a model fit. Yellow areas indicate the best possible fit given the input data and blue areas demonstrate the worst fits.*

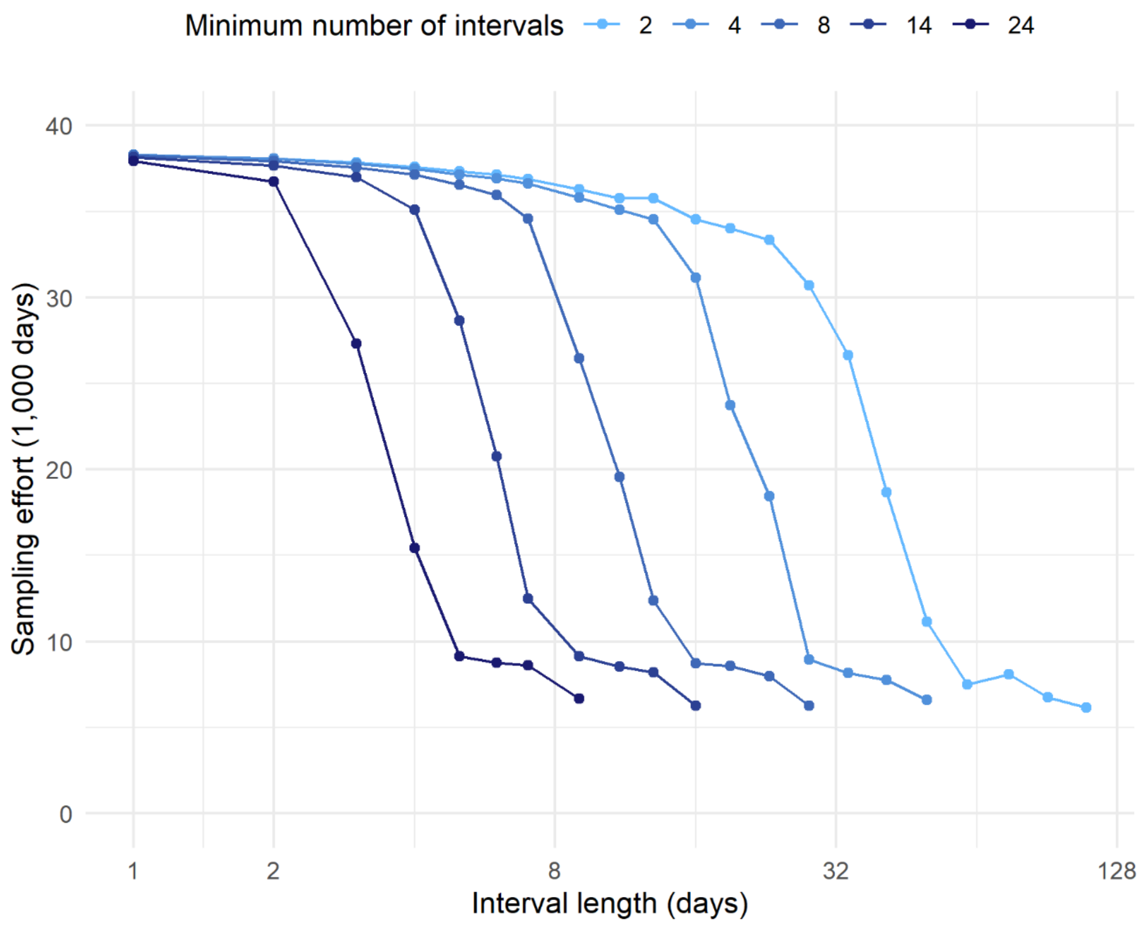


***Figure S4:*** *Sampling effort (in 1,000 days) of the camera trapping data of Zwerts et al. (2024) per interval length (in days) and minimum number of intervals needed per camera (nT). All cameras with a shorter individual sampling effort than dT x nT were disregarded.*

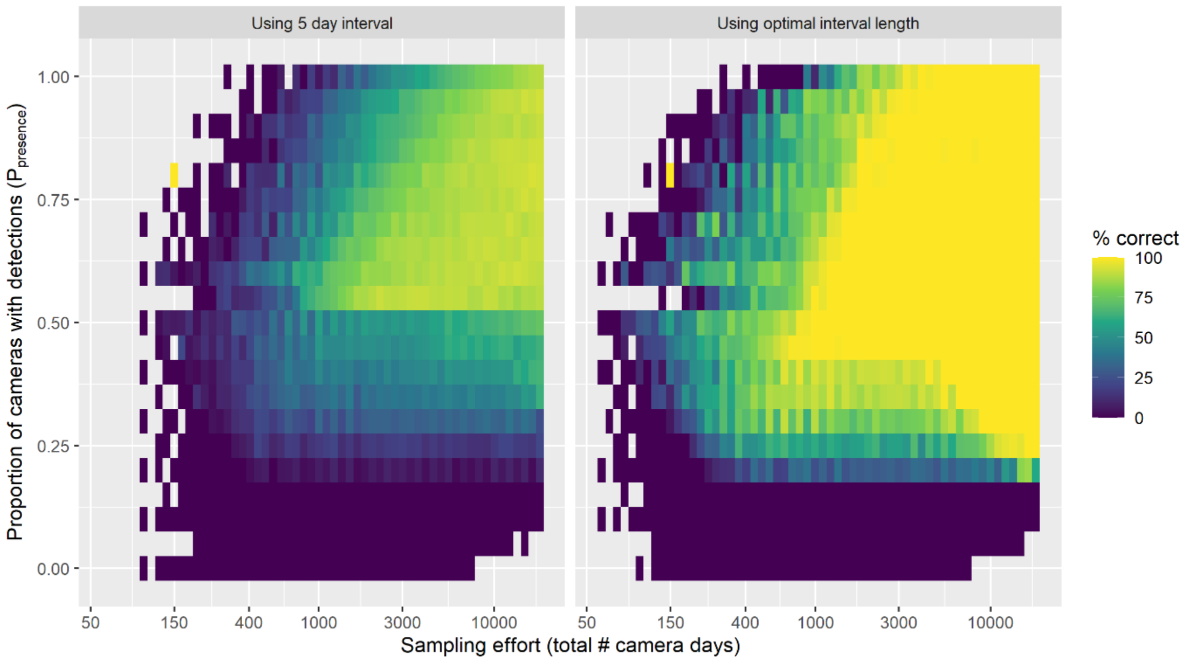


***Figure S5:*** *The percentage of instances in which the RN model provided correct results, using a 5 day interval (left panel) or the interval length that results in the best fitting RN model (right panel). Data was grouped per cluster of sampling effort (= total # camera days, x-axis) and proportion of cameras with detections ($P_{presence}$, y-axis).*
